# Stomatal and physiological response of contrasting *Z. mauritiana* (Lamk.) clones to water stress

**DOI:** 10.1101/077297

**Authors:** M. Kulkarni

**Affiliations:** A. Katz Department of Dryland Biotechnologies, The Jacob Blaustein Institutes for Desert Research (BIDR), Ben-Gurion University of the Negev (BGU), Sede-Boqer Campus, 84990 Sede-Boqer, Israel

**Keywords:** chlorophyll fluorescence, relative water content, leaf temperature, *Ziziphus*, water deficit

## Abstract

Water stress is one of the major limitations to fruit production worldwide. Identifying suitable indicators, screening techniques and quantifiable traits would facilitate the genetic improvement process for water stress tolerance. In the present study, we evaluated the ability of physiological parameters (Transpiration, E; *F*v/*F*m; leaf water ^potential, ψ^leaf^; leaf temperature, LT; and, leaf relative water content, RWC) to^ distinguish between contrasting *Z. mauritiana* clones subjected to a 30-d drought cycle. Four field-grown clones Seb and Gola (tetraploid) and Q 29 and B 5/4 (diploid) were ^studied. By 30 d after the onset of water stress treatment, the E, *F*v/*F*m, ψ^leaf ^and RWC^ of drought-stressed plants had declined significantly in all genotypes compared to values of well-watered treatments. However, the reductions were more severe in leaves of diploid clones. Under drought stress, the Seb and Gola, maintained higher E (31.5%), ^*F*v/*F*m (6.28%), ψ^leaf^; (11.2%), and RWC (9.3 %) than Q 29 and B 5/4 clones. In^ general, LT of drought-stressed plants was higher (~4°C) than that of well-watered plants but the relative increase was greater among later than former ones. Under maximum drought stress, LT of Seb and Gola clones was on average 3.0°C lower than that of Q 29 and B 5/4. Former clones yielded 20% more than later ones, mainly reason being (14.8%) less fruit drop as an effect of water stress. The results indicate that presented parameters can be reliable in screening for water stress tolerance ability, with ^*F*v/*F*m, ψ^leaf^, RWC and LT having the added advantage of being easily and quickly^ assessed.

## Introduction

Ber (*Ziziphus mauritiana* (Lamk.), Rhamnaceae is an important fruit crop in semi-arid tropics. *Z. mauritiana* (Lamk.) is a fruit tree species native to the semiarid regions of India, Pakistan, Australia, and South Africa. Efforts are being made to introduce this important fruit crop in Israel to diversify fruit production and provide alternative crop to farmers (Mizrahi and Nerd, 1996). Identifying suitable cultivars, growing conditions under different agro climatic conditions is first step towards commercialization of newly introduced fruit crops. Being arid zone, fruit growers in Israel relies heavily on irrigation to meet production goals. As, water for irrigation is a limited resource, its efficient utilization is critical in reducing production costs and for sustainable production. Identification of water stress tolerant clones is thus crucial for commercial production in arid zones of Negev desert in Israel where water supply is limited.

In *Z. mauritiana*, four distinct phonological phases have been characterized, namely vegetative growth (June to August, flowering (September to October) and fruiting (October to December) winter dormancy (February to May), under Negev desert conditions (Kulkarni et al., 2008a). Flowering and fruit growth period is critical water demand period (Katerji et al., 1993). Water relations and photosynthetic responses to water stress during this growth stage could therefore be useful in identifying better yielding clones with higher level of tolerance to water stress conditions.

Water stress adversely affects physiological processes such as stomatal conductance, leaf water potential (LWP), relative water content (RWC), leaf temperature (LT), transpiration(E), electron transport, photosynthesis and respiration which ultimately determine yield (Qing et al., 2001; Brodribb et al., 2003). A close association between photosynthesis, dry matter production and yield is reported with amount of water used by plant (Qing et al., 2001, Manoj and Swati, 2008). Genotypic variation for yield in species exists under water stress conditions (Aguilera et al., 1999). Therefore, ability of maintaining key physiological processes, such as photosynthetic ability during water stress conditions indicates potential for sustained productive ability of genotype.

Rong-hua et al. (2006), working with barley, showed that indirect and faster methods of measuring photosynthetic activity, such as chlorophyll a fluorescence technique, particularly the maximum photochemical efficiency of photosystem II – PSII (which can be accessed via the variable-to-maximum chlorophyll a fluorescence ratio, Fv/Fm), can be as effective as the more time-consuming gas exchange techniques in revealing differences between drought tolerant and susceptible genotypes. Other physiological parameters such as LT and relative water content (RWC) are also very responsive to drought stress and have been shown to correlate well with drought tolerance (Jamaux et al., 1997; Altinkut et al., 2001; Colom and Vazzana, 2003).

Different species uses different mechanisms for drought tolerance so it is essential to check reliability of earlier reported parameters between tolerant and susceptible clones in species under consideration (Colom and Vazzana, 2003; O’Neill et al., 2006; Rong-hua et al., 2006). To our knowledge, no studies have evaluated these relationships in these popular clones of *Z. mauritiana*. Careful selection of suitable physiological traits and quantifying them would be very valuable in selecting clones with better water stress tolerance. In the present study, we evaluated the ability of physiological parameters, namely the E, Fv/Fm ratio, LT, LWP and RWC to distinguish between four *Ziziphus mauritiana* clones for their water stress response.

## Materials and methods

This study was conducted in Sede Boker (longitude 34.78 °N; latitude 30.78 °E; altitude 450 m; mean annual rainfall 91 mm), Negev Desert, Israel, during the 2007-2008 in an experimental field with a sandy clay loam soil type. The experiment was arranged in a complete block design within a three-factor factorial, where the first factor was composed of four *Z. mauritiana* clones; the second factor was composed by two irrigation levels (wet and dry), and the third factor composed of three evaluation dates (0, 15 and 30 d after water deficit imposition), with four replicates. Data were collected from 6 trees per treatment per replication.

The four *Z. mauritiana* clones analyzed in this study were Seb, Gola, Q 29 and B 5/4. Seb and Gola are moderately vigorous, with small (average size 14.3±1.1 sq.cm) and thick leaves (average thickness 261±13µm) whereas Q 29 and B 5/4 had vigorous growth, with big (average size 25.6±1.8 sq.cm) and comparatively thinner leaves (average thickness 234±19 µm). Each clone was grafted on clonally propagated rootstock (*Z. nummularia*) planted in two rows, 3 m long, and 1.5 m apart in June, 2005. Chemical fertigation (Poly-Feed DRIP 23:7:23+2MgO with micronutrients, Haifa Chemicals Ltd.) was supplied through a drip system. On regular basis, plants were irrigated 56 L during the hot season (June to November) and 16 L during the cold wet season (December to March)

Two irrigation treatments [well-watered (+W) and water stress (-W)] were initiated in 26^th^ October, 2007 to 25^th^ November, 2007, during the flowering and fruiting period. The well-watered plots were irrigated at rate of 64 L/week, whereas in water stressed plots irrigation was stopped completely for 30 days. Soil moisture depletion was monitored periodically by monitoring % soil moisture on dry weight basis.

Flow cytometry analysis was performed as described by Garcia et al., (2008). *Pisum sativum* cv. Citrad was used as standard genome size reported to be 9.39 pg by Johanson et al. 1999. The absolute DNA content of samples was calculated based on values of M_1_ peak means:

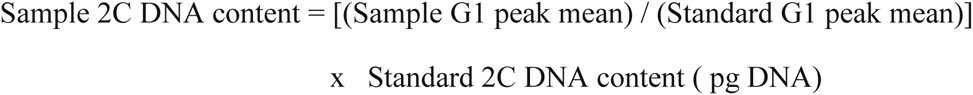

–as per Dolezel and Bartos (2005). *Z. mauritiana* (cv. Seb) [2n=48] was reported to be tetraploid (Mehetre and Dahat 2000) thus was used as internal standard to estimate ploidy level of rest of clones.

Stomatal observations were taken with an Axiomagera 1 LED (Zeiss) microscope. Length and width of fully opened stomata were measured using AxioVision Release 4.6.3.0 software (Carl Zeiss Imaging Solutions GmbH). Stomatal density was calculated by counting the number of stomata on the leaf epidermis in 1 mm^2^ microscopic field. Stomatal pore area (SPA) was calculated using pore length and pore width measurements. Four mature, fully expanded leaves located at the canopy’s periphery at a height of about 1.5 m above ground, exposed to full sunlight per tree from all trees under evaluation were used for this analysis.

Physiological parameters were measured three times during the study: at 0, 15 and 30 d after the onset of irrigation treatments (DAT) on cloudless days and between approximately 1100 h and 1500 h. Leaf transpiration rate (m mol H_2_0 m^−2^ s^−1^) was measured with a LI-1600 Porometer (LI-COR, Nebraska, USA) on five mature leaves per tree at 12:30-14:00 Hrs. Conversion of unit from μg H_2_0 m^−2^ s^−1^ to m mol H_2_0 m^−2^ s^−1^ was done as given by Nobel, 2005. During each measurement date, at least five leaves per tree were dark-adapted for 30 min using leaf clips (FL-DC, Opti-Science) before fluorescence measurements. The Fv/Fm ratio parameter was determined following the procedures of Maxwell and Johnson (2000), and used as to quantify the degree of drought-induced photoinhibition. Leaf temperature readings were collected using a hand-held infrared thermometer (Model OS530HR, Omega Engineering Inc., Stamford CT, USA) with leaf emissivity set at 0.95. During each LT measurement, the natural leaf orientation with respect to the sun was maintained to avoid shade effects.

Leaves used for E, Fv/Fm and LT were used for measurement of mid-day leaf water potential (Ψ_leaf_) and relative water content (RWC). Midday Ψ leaf was measured using a pressure chamber (SOIL Moisture Corporation, Santa Barbara, CA, USA) after Scholander et al. (1965) between 12:00 to 13:00 hrs for fully expanded outer canopy leaves (five per tree). Our main aim was to check maximum effect of water stress on plant water status so only mid-day water potential values were measured. To minimize errors due to water loss during procedure, leaves were cut and transported in humified polyethylene bags, and the base of chamber lines with moistened tissue. Leaf disks (1.3 cm diameter each) were collected with a cork borer and used to determine leaf relative water content (RWC) following the method of Matin et al. (1989). Five disks per tree were collected, immediately sealed in glass vials and quickly transported to the laboratory in an ice-cooled chest. Leaf disk fresh weights were determined within 2 h after excision. The turgid weight was obtained after rehydration in deionized water for 24 h at room temperature. After rehydration, leaves were quickly and carefully blotted dry with lint-free tissue paper before determining turgid weight. Dry weights were recorded after oven-drying leaf samples for 48 h at 80ºC.

After water stress treatment fruit yield was recorded as number of fruits per plant, average fruit weight and Yield in Kg per tree was for all treatments.

## Statistical analysis

Analysis of variance (ANOVA) was performed with JMP software (release 5.0.1a, SAS Institute Inc.) to determine significant differences (at p ≤05) between clones for all measured parameters. Tukey HSD test was used to separate means. Differences between cultivars for fruit yield per plant, number of fruits per plant, stomatal density and stomatal pore area (SPA) were determined using one-way ANOVA, with cultivar as the main effect and each parameter as a variable. All physiological parameters were analyzed by multivariate analysis considering clones and irrigation treatments as fixed effects and replication random effects. Evaluation dates were repeated observations in the analysis. Linear correlation analysis was used to determine the association among E, Fv/Fm, leaf RWC, SPA and fruit yield per tree.

## Results and discussion

Average air temperatures during the study period (September 2007 to October 2007) ranged from ca. 17.5 to ca. 29.0°C (Figure 1) and cumulative relative humidity was 70%. There was no occurrence of rainfall during all experiment. Soli relative water content decreased significantly up to 57.4 after 15 day drought and 42.3 after 30 days of imposing water stress as compared to well-watered condition (P≤0.001). The irrigation treatment received additional 64 L of water per week and water stressed plants utilized only soil water which went down slowly with increasing time period. As plants were only two years old possibility of roots to reach up to deeper moisture sources in soil are negligible.

**Figure 1:**
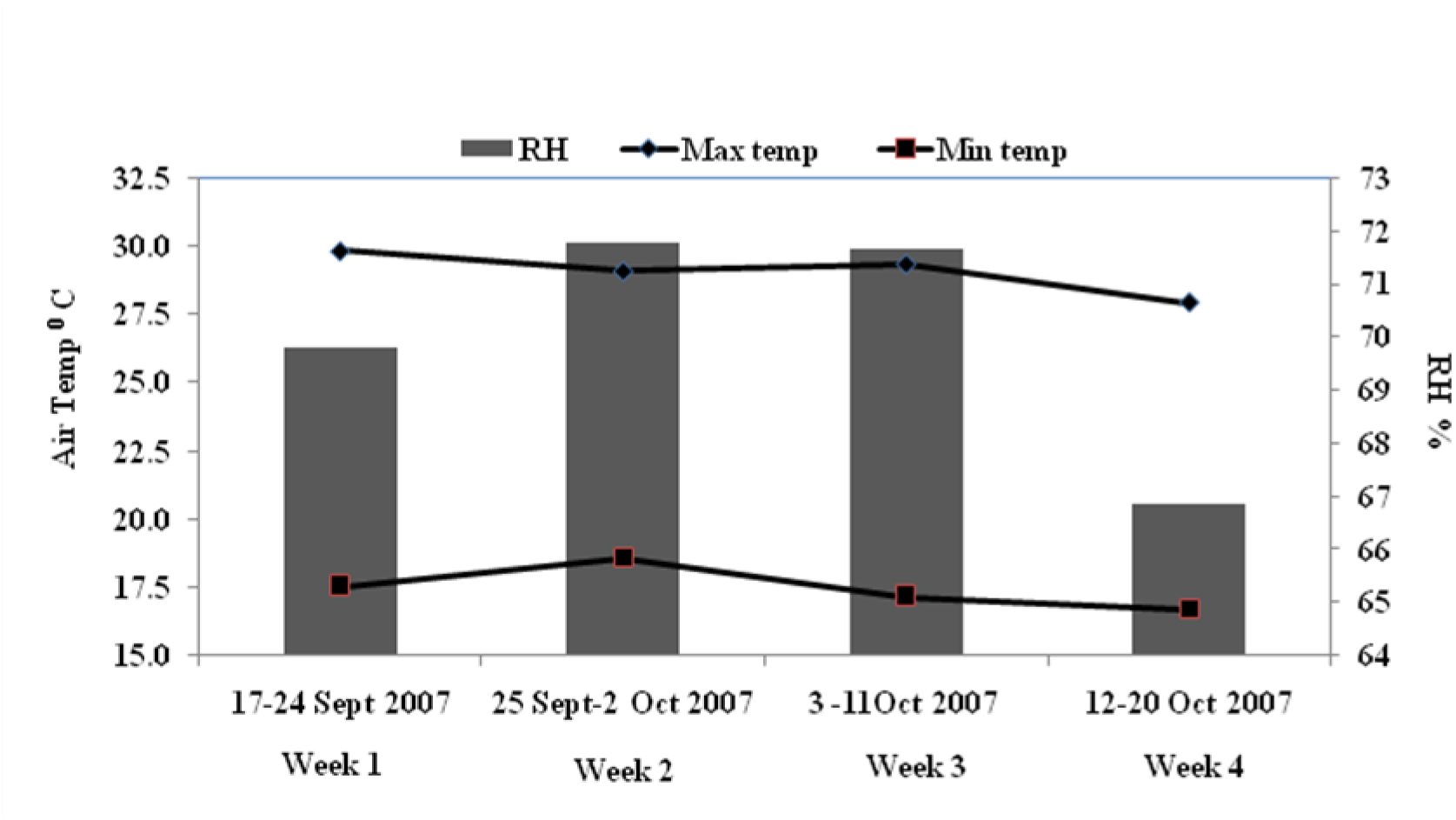
Weekly maximum, minimum air temperature and relative humidity during experimental period (September 2007 to October 2007). There was no occurrence of rainfall during experiment period.

### Ploidy level analysis

One dominant peak in all clones was obtained using flow cytometry analysis (FACS), indicated their relative nuclear DNA content (Fig.2). *Z. mauritiana* clones Seb and Gola samples were with significantly higher (2.19 – 2.35 pg) 2C nuclear DNA content than Q 29 and B 5/4 (1.70 – 1.79 pg) as shown in Table 1. These results indicated projected ploidy level of Seb and Gola to be tetraploid whereas Q 29 and B 5/4 being diploid 2C nuclear DNA content. These results supports our preliminary field study indicating Seb and Gola exhibiting better water stress tolerance under field condition than Q 29 and B 5/4 by maintaining better growth and vigor. Higher ploidy level with improved water stress tolerance is reported in many plant species (Stutz, 1989; Nutli and Zoblo, 2008; Kulkarni et al., 2008b).

**Table 1.** Genome size and expected ploidy based on FACS analysis

| Clone | Genome size (pg) | Expected ploidy |
| --- | --- | --- |
| Seb | 2.35 a | Tetraploid |
| Gola | 2.19 a | Tetraploid |
| Q 29 | 1.70 b | Diploid |
| B 5/4 | 1.79 b | Diploid |

**Figure 2:**
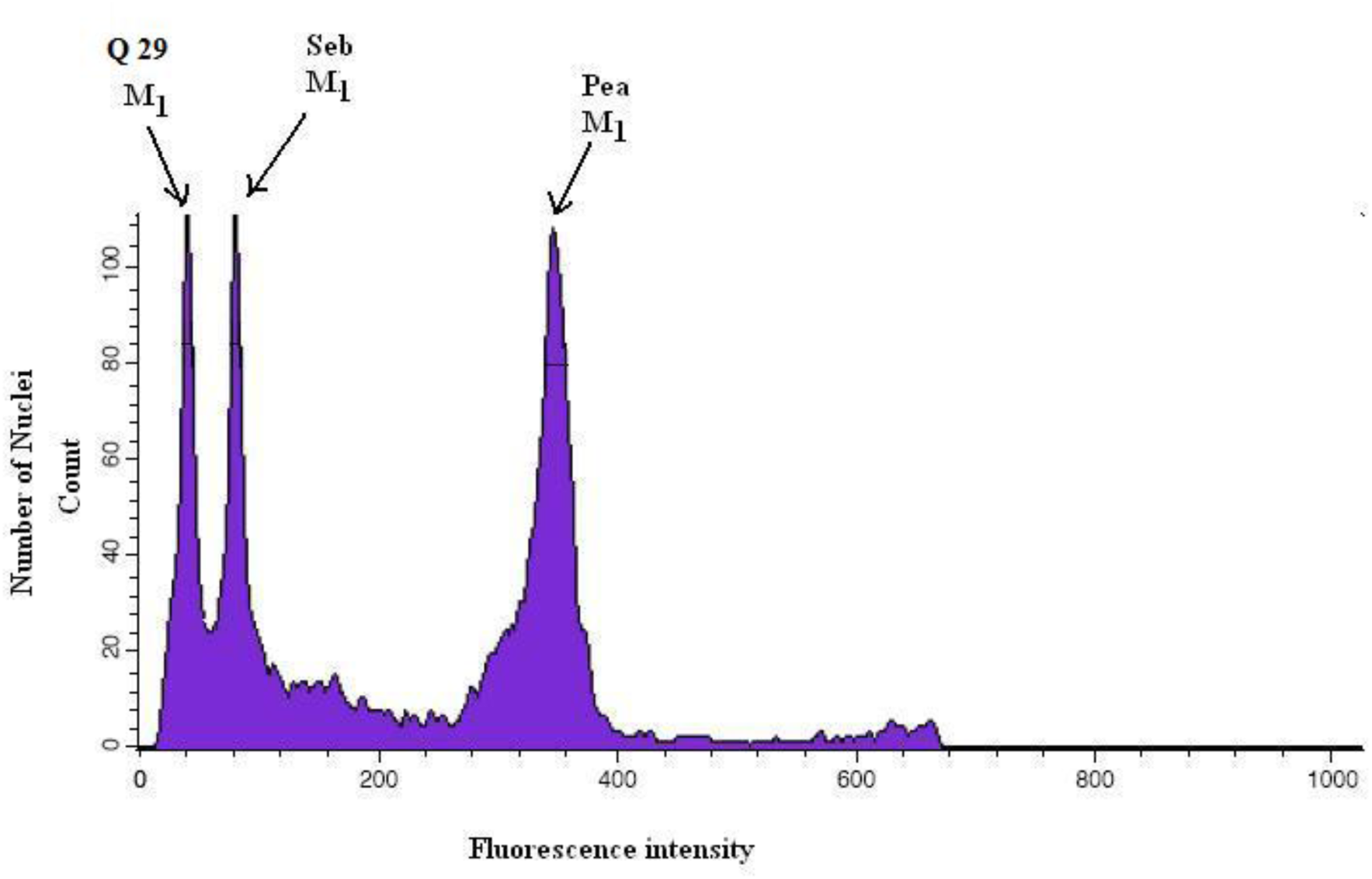
Representative flow cytometric histogram of nuclei isolated from leaves of *Z.mauritiana* clones. Seb clone had M_1_ peak at gate 80 whereas Q 29 had M_1_ peak at gate 45. Clone Seb used as internal standard being known tetraploid and known standard for nuclear DNA content in pg [Pea (*Pisum sativum* cv. Citrad)].

### Fruit yield under water stress

According to Kumar (2005), a plant or a group of plants showing better growth and productivity with limited soil moisture than other plants in a given set of similar environments is understood to be tolerant to water stress. Bearing this definition in mind and based on the last two years field performance under water stress conditions in south of Negev, the genotypes Seb and Gola were considered water stress tolerant. However, as Figure 3 shows, in the current experiment fruit yield for Seb and Gola differed significantly under well-watered (6.54 and 5.43 kg plant^−1^ respectievly) and drought-stress conditions (4.91 and 3.90 kg plant^−1^ respectievly), although the clones were able to maintain relatively high yield under both watering conditions. The genotypes Q 29 and B 5/4 could be considered water stress susceptible as fruit yield significantly decreased in response to water deficit application (Figure 3 A). Interestingly, clone B 5/4 was high yielding (at par with Seb) under irrigated condition but significantly higher reduction was observed (44.9%) under water stress as compared to rest of the clones. Main reason for reduction in yield was observed to be significantly high fruit drop in susceptible clones (Fig 3[B]; P≤ 0.01). Significantly lower fruit drop under water stress as compared to watered condition was observed in Seb (12.4%) and Gola(16.2%) whereas fruit drop was higher in clones Q29 (32.3%) and B 5/4 (23.5%). Overall, average reduction in fruit weight was observed to be (19.8%) but main reason for yield reduction under water stress being fruit drop.

**Figure 3:**
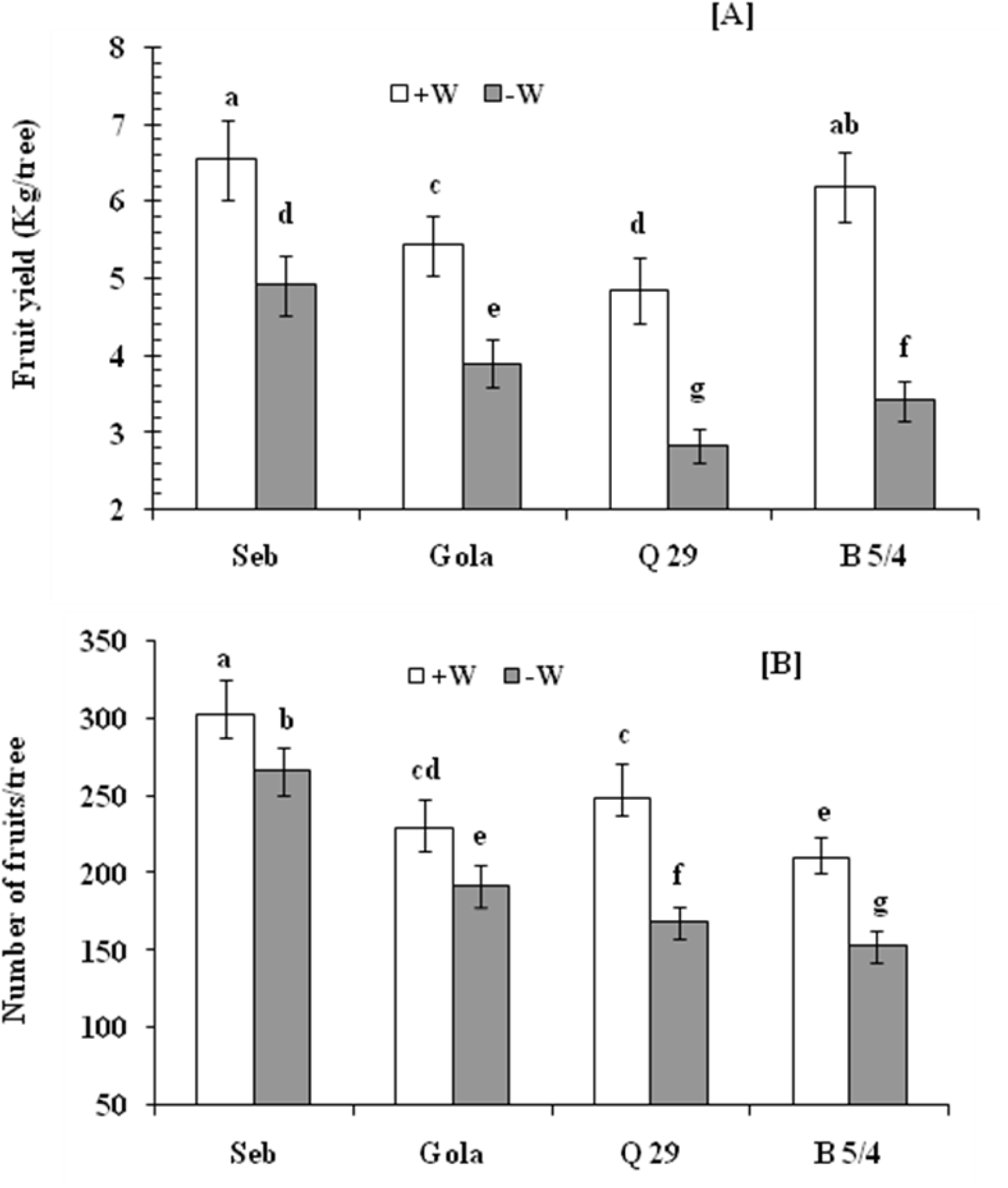
[A] Fruit yield (Kg/tree) and [B] number of fruits/ tree under normal and water stress condition in four clones of *Z. mauritiana*. Bars = SE. Different letters indicate significant differences by a Tukey HSD test, with P ≤0.05.

### Stomatal features

It was interesting to note that Seb and Gola (tetraploid clones) had significantly lower stomatal density on abaxial leaf surface per square mm (average 739.5±68) as compared to Q 29 and B 5/4 [diploid clones (average 1081±93)] under control condition. Under water stress condition, stomatal density increased significantly in all clones. Increase in stomatal density was significantly higher in clones Q 29 and B 5/4 as compared to Seb and Gola (Table 2). Early reports showed an increase in stomatal density and a decrease in cell size under water deficit, indicating that an adaptation to drought could occur (Wang and Gao, 2003; Yang et al., 2007; Gazanchian et al.,2007; Martinez et al., 2007).

**Table 2.** Stomata size, density and pore character variations on abaxial leaf epidermis in *Z. mauritiana* clones.

| Stomatal pore area ($\mu\text{m}^2$) | | Density (No.mm-2) | | Clone |
| --- | --- | --- | --- | --- |
| -W | +W | -W | +W |  |
| 75.8±7.1d | 139.4±12.6 a | 835±98b | 665±32 a | Seb |
| 63.2±5.3e | 107.4±9.9b | 1056±91cd | 814±60 b | Gola |
| 45.6±3.9f | 71.8±7.3 d | 1368±112g | 1164±27e | Q-29 |
| 42.6±4.1f | 81.3±6.2 c | 1243±105f | 998±24 c | B-5/4 |

Different letters indicate significant differences between cultivars at P≤0.05. (n=100 ± SE).

Lower stomatal density in drought tolerant rootstock (*Z. nummularia*) as compared to *Z. mauritiana* is reported by Bankar and Prasad, 1992. Overall negative relation between stomatal size and density was observed in four clones under study. Bigger stomatal size and lower density was related to ploidy level in *Z. jujube* (Gu et al., 2005). Stomatal density was also negatively correlated with stomatal size under different water conditions in *Z. jujube* leaves (Zheng et al., 2006) and *Platanus acerifolia* leaves (Zhang et al., 2004). Nevertheless, different effects of abiotic factors on stomatal size may depend on plant species/varieties (Maherali et al., 2002; Liu et al., 2006). There was significant effect of water stress on stomatal pore area in all clones under study but this effect was with higher magnitude on Q 29 and B 5/4 as compared to Seb and Gola (Table 2; Fig.4). Vincent et al. (2005) reported similar stomatal regulation in drought-resistant *Cocos nucifera*: stomata were partially closed in tolerant tall cultivars and completely closed in sensitive dwarf cultivars.

**Figure 4:**
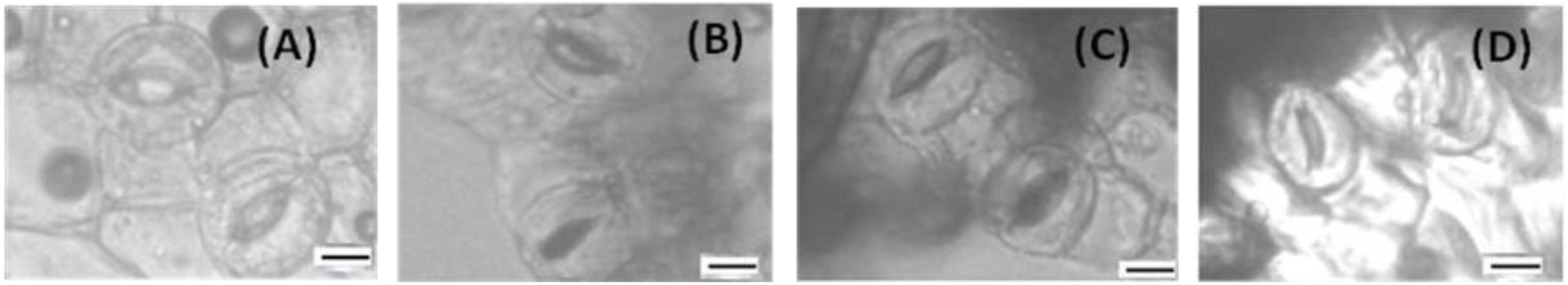
Effect of water stress on abaxial stomata: (A) Seb and (B) Q 29 under normal irrigation condition; (C) Seb (partially closed) and (D) Q 29 (Completely closed) under water stress conditions (Scale bar = 5 µm).

### Physiological indicators of water stress response

Significant genotype by irrigation (GxW), genotype by evaluation date (GxD) and irrigation by evaluation date (WxD) interactions were observed for photosystem II (PSII) photochemical efficiency (Fv/Fm) measurements (Table 3). Under well-watered conditions, tolerant as well as susceptible genotypes maintained high Fv/Fm values (~0.76-0.77). Drought stress generally resulted in decreased Fv/Fm in all clones, but proportionate decrease was more in diploid clones (Tables 4 and 5). PSII activity reduction as an effect of photoinhibition due to water stress is indicated by Fv/Fm reduction (Maxwell and Johnson, 2000). The ability to maintain higher Fv/Fm by Seb and Gola under drought stress thus indicates a high efficiency of radiation use possibly for photochemistry and carbon assimilation. Maintaining higher value of Fv/Fm under water stress associated with water stress tolerance and reduction in these values associated with susceptibility are reported in recent publications (Percivel and Sheriffs, 2002; Raveh et al., 1995; Yang et al., 2005; Colom and Vazzana, 2003). Clifford et al., 1998 did not found significant difference for Fv/Fm value between normal and water stressed *Z. mauritiana* plants during 13 day water stress experiment. But plants are greater sink at flowering and fruiting phase which amplifies water stress effect could be assumed to result in significant reduction in Fv/Fm values indicating sever water stress under presented experimental conditions.

**Table 3.**
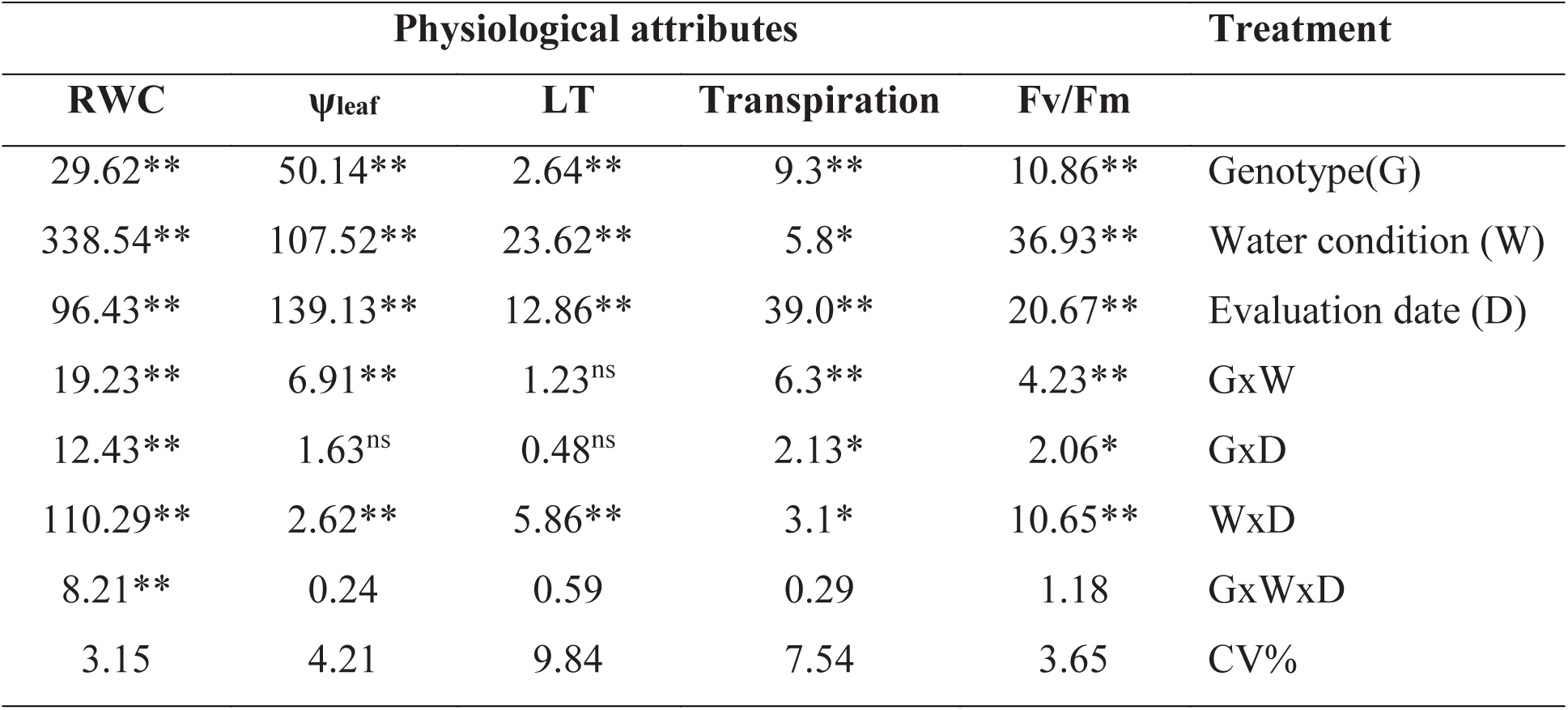
Analysis of variance (*F* values) for PSII photochemical efficiency (*F*v/*F*m), transpiration (E), leaf temperature (LT), midday leaf water potential (ψ_leaf_) and leaf relative water content (RWC) of four *Z. mauritiana* clones grown under well-watered and drought conditions and measured on three dates. * *P* < 0.05; ** *P* < 0.01.

| Physiological attributes |  |  |  |  | Treatment |
| --- | --- | --- | --- | --- | --- |
| RWC | $\psi_{\text{leaf}}$ | LT | Transpiration | $F_v/F_m$ | |
| 29.62** | 50.14** | 2.64** | 9.3** | 10.86** | Genotype(G) |
| 338.54** | 107.52** | 23.62** | 5.8* | 36.93** | Water condition (W) |
| 96.43** | 139.13** | 12.86** | 39.0** | 20.67** | Evaluation date (D) |
| 19.23** | 6.91** | 1.23<sup>ns</sup> | 6.3** | 4.23** | GxW |
| 12.43** | 1.63<sup>ns</sup> | 0.48<sup>ns</sup> | 2.13* | 2.06* | GxD |
| 110.29** | 2.62** | 5.86** | 3.1* | 10.65** | WxD |
| 8.21** | 0.24 | 0.59 | 0.29 | 1.18 | GxWxD |
| 3.15 | 4.21 | 9.84 | 7.54 | 3.65 | CV% |

**Table 4.** PSII photochemical efficiency (Fv/Fm), Transpiration (E), leaf temperature (LT), Leaf water potential (ψ_leaf_) and leaf relative water content (RWC) of four *Z. mauritiana* clones grown under well-watered (+W) and water stress (-W) conditions. Measurements were taken at 15 days after the onset of irrigation treatments. Means for tolerant and susceptible clones in a same row and within a same attribute column and having the same letter are not significantly different at 0.05 probability level (Tukey’s test).

| Physiological attributes |  |  |  |  |  |  |  |  |  | Clone |
| --- | --- | --- | --- | --- | --- | --- | --- | --- | --- | --- |
| RWC | | $\psi_{\text{leaf}}$ | | LT | | E | | Fv/Fm | | |
| -W | +W | -W | +W | -W | +W | -W | +W | -W | +W |  |
| 81.3 b | 88.1 a | -2.15 c | -1.29 b | 32.0b | 29.9a | 3.1 d | 3.9 c | 0.72c | 0.76a | Seb |
| 79.4 b | 88.2 a | -2.33 d | -1.46 a | 32.5b | 28.7a | 2.9 e | 5.3 b | 0.71c | 0.77a | Gola |
| 76.8 c | 88.3 a | -2.52 e | -1.34 b | 34.8c | 27.6a | 3.2 d | 6.0 a | 0.71c | 0.77a | Q 29 |
| 77.1 c | 87.2 a | -2.48 e | -1.45 a | 35.1c | 28.3a | 2.6 f | 5.2 b | 0.69d | 0.74b | B 5/4 |

**Table 5.** PSII photochemical efficiency (Fv/Fm), Transpiration (E), leaf temperature (LT), Leaf water potential (ψ_leaf_) and leaf relative water content (RWC) of four *Z. mauritiana* clones grown under well-watered (+W) and water stress (-W) conditions. Measurements were taken at 30 days after the onset of irrigation treatments. Means for tolerant and susceptible clones in a same row and within a same attribute column and having the same letter are not significantly different at 0.05 probability level (Tukey’s test).

| Physiological attributes |  |  |  |  |  |  |  |  |  | Clone |
| --- | --- | --- | --- | --- | --- | --- | --- | --- | --- | --- |
| RWC | | $\psi_{\text{leaf}}$ | | LT | | E | | Fv/Fm | | |
| -W | +W | -W | +W | -W | +W | -W | +W | -W | +W |  |
| 78.7b | 87.1a | -2.39c | -1.27b | 33.6b | 29.6a | 2.6 d | 4.2 c | 0.69c | 0.78a | Seb |
| 76.3b | 86.9a | -2.52d | -1.42a | 34.3b | 29.1a | 2.4 d | 5.0 b | 0.68c | 0.78a | Gola |
| 71.4c | 87.3a | -2.62e | -1.36b | 37.1d | 27.4a | 2.0 e | 6.1 a | 0.64d | 0.77a | Q 29 |
| 70.3c | 87.2a | -2.69e | -1.43a | 36.8c | 28.1a | 1.8 e | 5.1 b | 0.65d | 0.76b | B 5/4 |

Transpiration responses to water stress were similar to those obtained for Fv/Fm with the exception that transpiration data varied more as compared to Fv/Fm. Transpiration was significantly affected by GxW and WxD interactions (Table 3). Transpiration declined progressively with exposure to drought but the decline was more severe in clones Q 29 and B 5/4, as could be deduced from Tables 4 and 5. Q 29 had the highest transpiration rate under normal conditions could be attributed to higher stomatal density (Table 2). Transpiration rate values under drought conditions were maintained at higher proportions as compared to control values by Seb and Gola (45.3%) but Q 29 and B 5/4 had the higher reductions (64.1%) as compared to respective control values at highest water stress level i.e. 30 days after imposing water stress (Tables 5). It is known that the maintenance of transpiration under conditions of water stress constitutes a mechanism of water stress tolerance, as has been shown for *Hevea brasiliensis*, *Psidium guajava* and *Anacardium occidentale* trees (Ozinaldo et al., 2007). Similar results were also reported for *Quercus coccifera, Ceratonia siliqua, Pistacia terebinthus, Olea oleaster* (Sakcali and Özturk, 2004) and *Hordeum vulgare* (Rong-hua et al., 2006).

Drought stress generally resulted in an overall increase in LT for all the clones, regardless of tolerance susceptibility classification (Tables 3, 4 and 5). However, LT of clones, Q 29 and B 5/4 responded sharper to drought stress (by 15 DAT) than that of Seb and Gola (Table 4). The highest average increase in LT was observed in Q 29 whereas the lowest average increase was recorded for Seb (Table 4). Overall, clones classified as susceptible generally had higher average LT readings (~36.9°C) under drought stress conditions than those classified as tolerant (~33.9°C; Table 5). The increase in LT was probably due to reduced evapotranspirational cooling, resulting from drought-induced stomatal closure (Fig.4). As stomata close in response to water deficit stress, transpirational cooling ceases, leading to a rise in leaf temperature (Luquet et al., 2003; Jones, 2004). While stomatal closure under increasing water stress helps to prevent development of lethal water deficits, but it can lead to lethal temperatures under high radiation and temperature conditions.

It was evident from partial stomatal closure in tolerant clones under water deficit conditions that they maintain better water status and have dynamic control over stomtatal regulation leading to sustained transpirational cooling. This mechanism leads to sustained CO_2_ influx towards chloroplasts, maintained photosynthesis rate and crop yield (Kumar, 2005). Although LT data varied more as compared to Fv/Fm, perhaps due to reflection of solar radiation during measurement and changing wind conditions, water stress induced differences between Seb, Gola (tetraploid) and Q29, B5/4 (diploid) clones were still apparent. Possible error due to excessive reflection of solar radiation during measurement may lead to overestimation of LT which could be avoided by temporary shading of leaf before measurement (Leigh et al. 2006).

### Trends in plant water relations during water stress

Relative water content (RWC) of leaves was determined along with measurements of leaf water potential in the repeated measurements. Similar leaf water potential values were obtained, which positively correlated with the RWC of the genotypes (Fig. 5, r^2^=0.93). Significant decrease observed in RWC as well as lowered LWP under water stress indicated level of water stress imposed after withholding irrigation.

**Figure 5:**
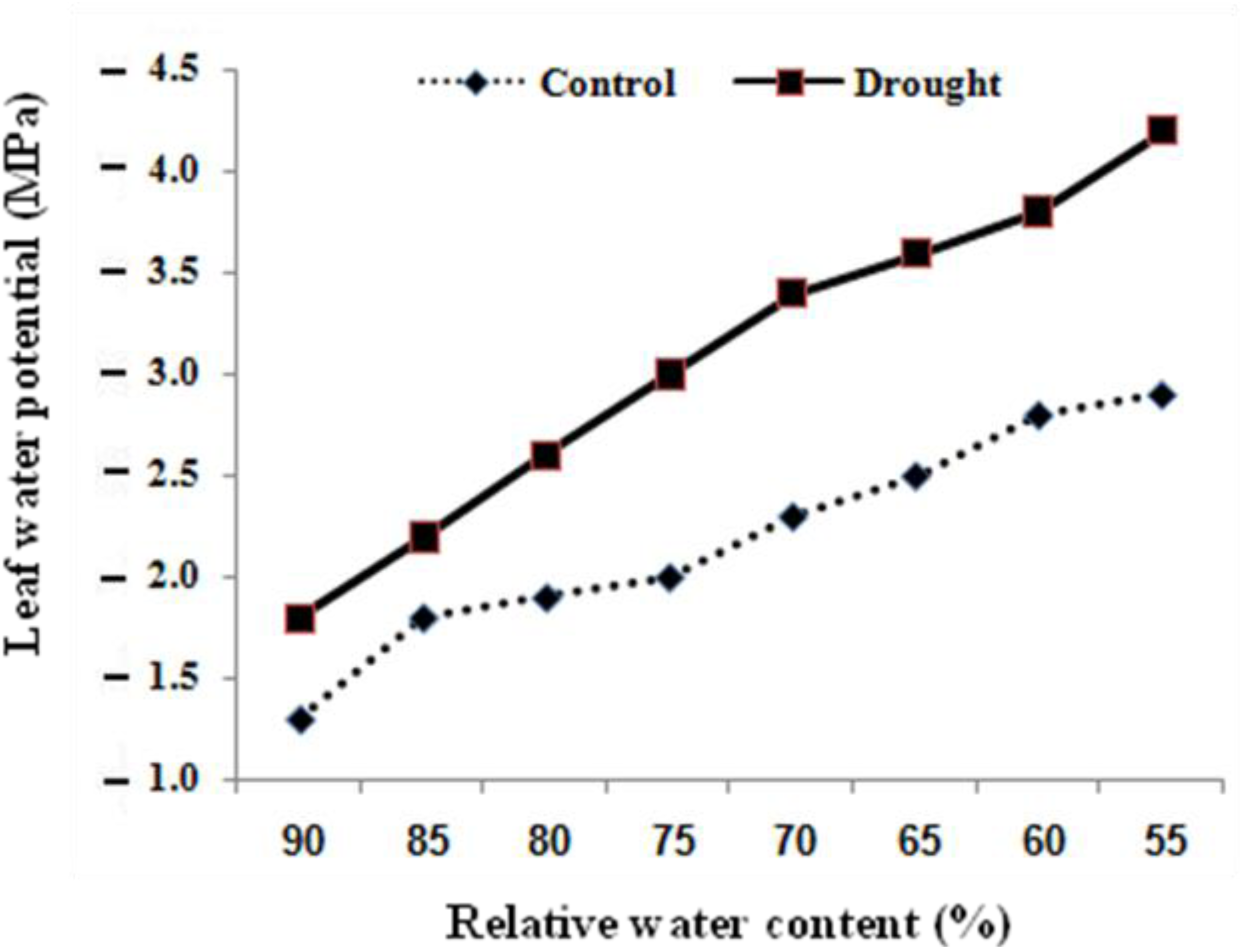
Relationship between leaf water potential and relative water content of leaves of well-watered (dotted line) and water-stressed plants (solid line) of *Ziziphus mauritiana* clones under study. (n =140, r^2^ = 0.93)

Leaf relative water content (RWC) was the only parameter for which the GxWxD interaction was significant (Table 1). At the onset of differential irrigation treatments RWC was similar (~88%) among all clones (Table 3) but water deficit stress resulted in progressive decline in RWC (Tables 4 and 5). By 15 DAT, significant differences in RWC had developed with clones from the tolerant tetraploid clones (Seb and Gola) maintaining a relatively higher average RWC (~80%) than those in the susceptible clones [(Q29 and B 5/4 (~76%, Table 4)]. This trend was more evident at 30 DAT (Table 5).

On average, water stress-induced reduction in RWC occurred to a greater extent in the water stress-susceptible clones Q 29 (19.3%) and B 5/4 (21.4%), and to a lesser degree in the tolerant Seb (9.7%). The average water stress-induced reduction in RWC was 12.9% for the tolerant clones and 20.3% for the susceptible clones. Clones in the tolerant category in this study had relatively high RWC values compared to those in the susceptible category thus confirming their empirical classification as water stress tolerant. Clifford et al., 1998 reported reduction of 14% RWC content after a period of 13 day water stress cycle to *Z. mauritiana* clones Seb and Gola. Physiological functioning and growth under water stress depends on degree of cell and tissue hydration for which RWC is an important indicator. Maintaining relatively higher RWC during water stress as an indicator of drought tolerance is reported in numerous earlier studies (Jamaux et al., 1997; Altinkut et al., 2001; Colom and Vazzana, 2003).

Significant genotype by irrigation (GxW) and irrigation by evaluation date (WxD) interactions were observed for mid-day LWP measurements (Table 3). Under well-watered conditions, tolerant as well as susceptible clones maintained lower LWP (~ −1.29 to −1.46 MPa). It was indicative that reduction in RWC was high enough to cause reduction in LWP. It could be assumed that water stress led to significant drop in LWP in transpiring leaves such that cells began to lose turgor, triggering stomatal closure (Brodribb et al., 2003).

Midday values of LWP in full irrigation were higher than the drought treatment for the entire experimental period (Table 4 and 5). Midday LWP started to decrease from 10 days after withholding irrigation reaching lowest values (MPa) of ~ −2.69 in Q 29 and B 5/4 whereas significantly higher values of LWP were observed under similar stress level (~ −2.45) in Seb and Gola. Significantly lowered values of midday LWP in former indicate higher magnitude of water stress as compared to later clones. Reduced midday leaf water potential from −1.1 to −2.4 MPa during a water stress period of 13 days was reported by Clifford et al., 1998, as in our case plants were at flowering and fruiting phase and duration of drought was almost double could impose a water stress of higher magnitude.

**Table 6.**
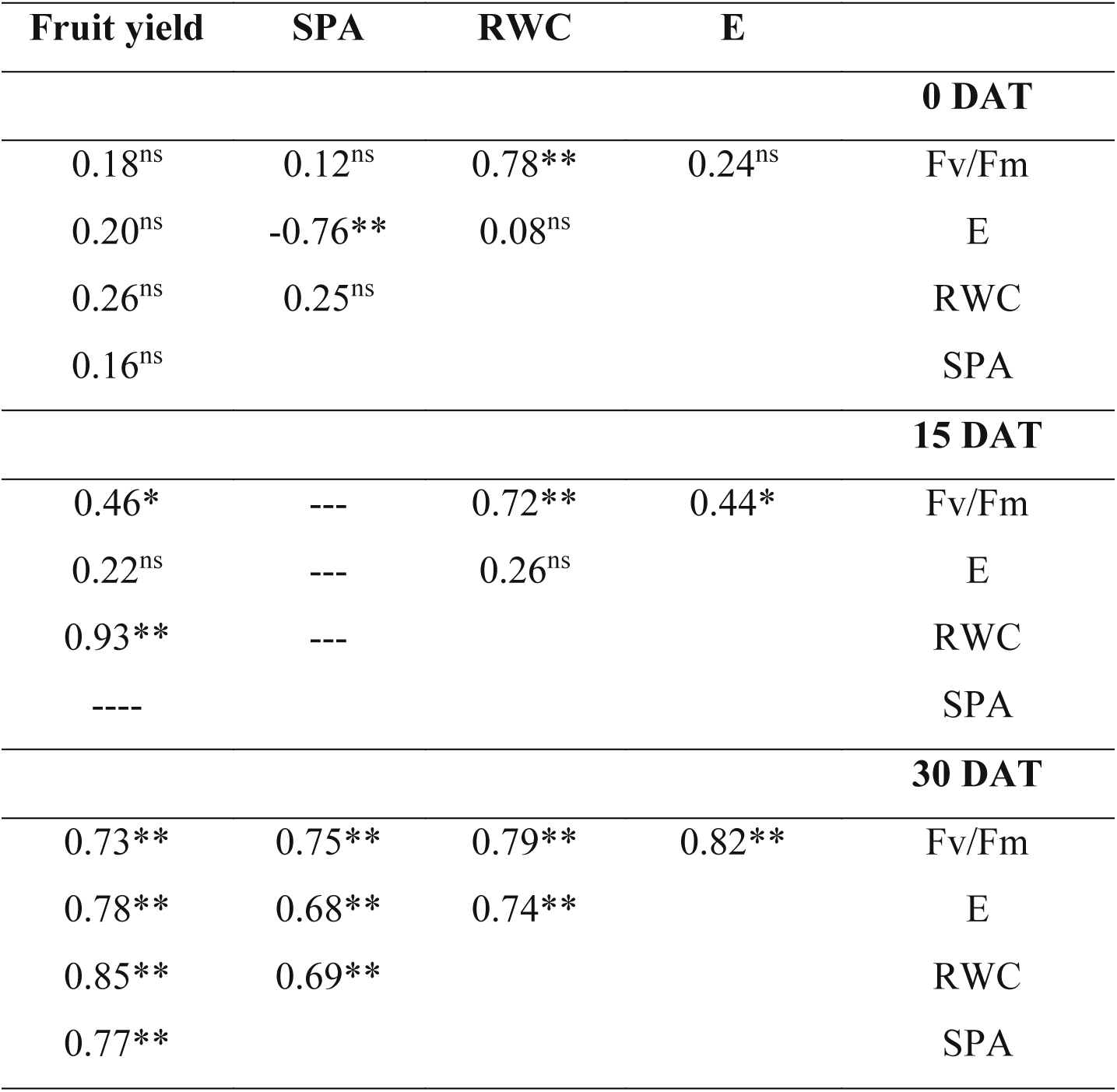
Pearson’s correlation coefficients for association among chlorophyll PSII photochemical efficiency (Fv/Fm), Transpiration (E), leaf relative water content (RWC), stomatal pore area (SPA) and fruit yield of four *Z. mauritiana* clones grown under two watering treatments measured on 0, 15 and 30 days after treatment imposition (DAT). ns, not significant; * P < 0.05; ** P < 0.01. (**Note:** stomatal pore area (SPA) was measured at initial and maximum water stress conditions only)

| **Fruit yield** | **SPA** | **RWC** | **E** |  |
| --- | --- | --- | --- | --- |
| **0 DAT** |  |  |  |  |
| 0.18<sup>ns</sup> | 0.12<sup>ns</sup> | 0.78** | 0.24<sup>ns</sup> | Fv/Fm |
| 0.20<sup>ns</sup> | -0.76** | 0.08<sup>ns</sup> |  | E |
| 0.26<sup>ns</sup> | 0.25<sup>ns</sup> |  |  | RWC |
| 0.16<sup>ns</sup> |  |  |  | SPA |
| **15 DAT** |  |  |  |  |
| 0.46* | --- | 0.72** | 0.44* | Fv/Fm |
| 0.22<sup>ns</sup> | --- | 0.26<sup>ns</sup> |  | E |
| 0.93** | --- |  |  | RWC |
| ---- |  |  |  | SPA |
| **30 DAT** |  |  |  |  |
| 0.73** | 0.75** | 0.79** | 0.82** | Fv/Fm |
| 0.78** | 0.68** | 0.74** |  | E |
| 0.85** | 0.69** |  |  | RWC |
| 0.77** |  |  |  | SPA |

Before the onset of treatments there was no significant correlation between the parameters, with the exception of a positive association between Fv/Fm and RWC, whereas a significant negative correlation was observed between E and SPA (Table 6). A modest, although significant, positive correlation was found among Fv/Fm, and E as well at 15 DAT as at 30 DAT. The degree of association amongst these parameters varied between 0.33 to 0.50 at 15 and 30 DAT, respectively.

On the other hand, stomatal pore area showed a significant negative correlation with E, at under normal irrigation treatment (−0.66 to −0.81). Under water stress conditions this correlation was found to be significant (0.68). It was interesting to note that, relation between Fv/Fm and SPA was non-significant under unstressed condition but was found significant under water stress conditions. These results indicate that stomatal pore area may be more closely related to photosynthetic character, and water deficit may enhance the correlations.

The fruit yield was found to be positively correlated with Fv/Fm, E, SPA and RWC during the whole evaluation period under drought conditions. These results demonstrated that all traits were affected mutually, positively, and were consistent in response to water deficit condition. Therefore, these parameters might be used as a selection criterion for fruit productivity in *Ziziphus* under water stress. This is in accordance to Lutfur et al., (1999), Rong-hua et al., (2006), Manoj and Swati, (2008), that worked with tomato, barley, hot pepper.

## Concluding remarks

Water stress is one of the major reasons to fruit productivity worldwide and a possible solution is to improve the water stress tolerance of fruit crops through genetic improvement for water stress tolerance. To achieve this goal, a set of reliable traits that can be easily screened is needed. Overall, the yield reduction after 30 days of water stress observed for the tolerant clones (Seb and Gola) in lower proportion and the considerable correlation with the parameters studied turns this approach quite promising. Although all the traits and techniques evaluated in this study were reliable in distinguishing between tolerant and susceptible *Ziziphus mauritiana* clones, Fv/Fm and RWC seem to be the most promising for water stress tolerance screening. As *Z. mauritiana* is relatively water stress tolerant fruit crop, the best responses for screening for water stress tolerance could be achieved after 30 d under water limitation during the flowering and fruiting phase.

## Acknowledgement

Author is thankful to the Albert Katz Department of Dryland Biotechnologies, Blaustein Institutes for Desert Research, Ben Gurion University of Negev for Post-doctoral fellowship. The author thank Dr. Noemi Tel-Zur; Prof. Yoseph Mizarhi for experimental plant material and Yossi Moyal’s help for water stress experiment.

